# Increased oxidative stress tolerance of a spontaneously-occurring *perR* gene mutation in *Streptococcus mutans* UA159

**DOI:** 10.1101/2020.09.24.312702

**Authors:** Jessica K. Kajfasz, Peter Zuber, Tridib Ganguly, Jacqueline Abranches, José A. Lemos

## Abstract

The ability of bacteria such as the dental pathogen *Streptococcus mutans* to coordinate a response against damage-inducing oxidants is a critical aspect of their pathogenicity. The oxidative stress regulator SpxA1 has been proven to be a major player in the ability of *S. mutans* to withstand both disulfide and peroxide stresses. While studying spontaneously-occurring variants of an *S. mutans* Δ*spxA1* strain, we serendipitously discovered that our *S. mutans* UA159 host strain bore a single nucleotide deletion within the coding region of*perR*, resulting in a premature truncation of the encoded protein. PerR is a metal-dependent transcriptional repressor that senses and responds to peroxide stress such that loss of PerR activity results in activation of oxidative stress responses. To determine the impact of loss of PerR regulation, we obtained a UA159 isolated bearing an intact *perR* copy and created a clean *perR* deletion mutant. Our findings indicate that loss of PerR activity results in a strain that is primed to tolerate oxidative stresses in the laboratory setting. Interestingly, RNA-Seq and targeted transcriptional expression analyses reveal that PerR has a minor contribution to the ability of *S. mutans* to orchestrate a transcriptional response to peroxide stress. Furthermore, we detected loss-of-function *perR* mutations in two other commonly used laboratory strains of *S. mutans* suggesting that this may be not be an uncommon occurrence. This report serves as a cautionary warning regarding the so-called domestication of laboratory strains and advocates for the implementation of more stringent strain authentication practices.

**Importance:** A resident of the human oral biofilm, *Streptococcus mutans* is one of the major bacterial pathogens associated with dental caries. This report highlights a spontaneously-occurring mutation within the laboratory strain *S. mutans* UA159, found in the coding region of *perR*, a gene encoding a transcriptional repressor associated with peroxide tolerance. Though *perR* mutant strains of *S. mutans* showed a distinct growth advantage and enhanced tolerance toward H_2_O_2_, a Δ*perR* deletion strain showed a small number of differentially expressed genes as compared to the parent strain, suggesting few direct regulatory targets. In addition to characterizing the role of PerR in *S. mutans*, our findings serve as a warning to laboratory researchers regarding bacterial adaptation to *in vitro* growth conditions.

## INTRODUCTION

Transition metals are essential micronutrients for cell life. However, when found in excess, these same metals can be extremely harmful, in part because of the deleterious effects of Fenton chemistry, by which intermingling of H_2_O_2_ and soluble ferrous iron (Fe^2+^) or copper (Cu^2+^) results in formation of the highly reactive hydroxyl radical (HO·) (1). To maintain metal homeostasis and avoid reactive oxygen species (ROS) intoxication, bacteria employ a number of metal-sensing regulators (metalloregulators) that are used to maintain metal homeostasis and coordinate oxidative stress responses. Among the most extensively studied metal-sensing regulators are the ferric uptake regulator (Fur) protein family (2). Members of this family function as homodimers and include sensors for iron (Fur), zinc (Zur), manganese (Mur), nickel (Nur), and peroxide (PerR). PerR is an oxidative stress regulator that senses peroxide stress when bound to a metal co-factor that, depending on the bacterial species, can be iron, manganese, or both (2, 3). PerR has been shown to function as a transcriptional repressor by binding directly to operator sites within the promoter region of a gene target (4–6). Among gram-positive bacteria, PerR is associated with repression of oxidative stress-detoxification genes such as *dpr* (encoding an iron-binding peroxide resistance protein), *sodA* (superoxide dismutase), and *ahpCF* (alkyl hydroperoxidase), though the composition of regulons governed by PerR varies among bacterial species and even between strains within a species (7–11). In general, the genes regulated by PerR can be divided into two major groups: genes directly involved in ROS detoxification, and genes involved in metal homeostasis (12). Of note, PerR homologs are widespread in Firmicutes but are also occasionally found in gram-negative bacteria (13–15).

Among Firmicutes, *Streptococcus mutans* is a resident of the dental biofilm shown to be a major contributor to dental caries development and, occasionally, associated with systemic infections such as infective endocarditis (16). The association of *S. mutans* with dental caries is strongly linked with its ability to tolerate environmental stresses encountered in its environmental niche, namely acidic and oxidative stresses (17). Multiple sources contribute to the oxidative stresses found in dental plaque, including the capacity of the oral microbial community as a whole to reduce oxygen into ROS, and the direct production of H_2_O_2_ by peroxigenic bacteria. The transcriptional regulators SpxA1 and SpxA2 have been proven to function as central regulators of the oxidative stress response of *S. mutans*, with SpxA1 serving as a direct activator of the genes classically associated with the oxidative stress response whereas SpxA2, whose major role has been recently linked to cell envelope homeostasis, serves as a backup system (18, 19). As part of our ongoing characterization of *S. mutans spx* mutant strains, we found that the *S. mutans* Δ*spxA1* strain was hypersensitive to arginine, a phenotype that was linked to alkaline stress tolerance (P. Zuber, unpublished data). As we observed the emergence of stable Δ*spxA1* phenotypic revertants on plates containing arginine, we subjected those revertants to whole genome sequencing (WGS) for the purpose of identifying the locus or loci linked to arginine tolerance. In addition to linking loss-of-function mutations in an iron transporter to the rescue of arginine tolerance in the Δ*spxA1* strain, we unexpectedly discovered that all isolates subjected to WGS, including the original arginine-sensitive Δ*spxA1* isolate, bore a mutation in the *perR* gene that likely renders PerR inactive. In light of this unexpected finding, we sought to characterize the impact of PerR on *S. mutans* physiology. Despite displaying a rather modest role in the transcriptional regulation of oxidative stress genes, we showed that loss of PerR regulation primed *S. mutans* for growth in the presence of oxidative stresses under *in vitro* conditions. *In silico* and PCR sequencing analyses of UA159 laboratory stocks as well as other widely used *S. mutans* strains revealed that the occurrence of spontaneous *perR* mutations may not be uncommon in laboratory strains.

## RESULTS AND DISCUSSION

### Loss-of-function mutations of the *smu995-998* iron transporter in the Δ*spxA1* strain leads to serendipitous identification of a spontaneous *perR* mutation that increases oxidative stress tolerance

The work described here began as an exploration of the initial observation that our Δ*spxA1* strain was hypersensitive to arginine stress. Though the Δ*spxA1* strain showed enhanced sensitivity to growth in the presence of arginine as compared to the parent strain, a number of revertant colonies were identified that regained the ability to grow in the presence of arginine (Fig. S1A). Four stable revertant isolates and the original Δ*spxA1* strain were subjected to WGS analysis to identify the locus in which the mutation(s) that restored arginine tolerance to the Δ*spxA1* strain resided. When compared to the annotated *S. mutans* UA159 genome (NC_004350.2), all four colony revertants (numbered R1, R4, R6 and R7) harbored single nucleotide polymorphisms (SNPs) in the *smu997* or *smu998* genes, which encode the ATP-binding and substrate-binding proteins of an iron transport system (Fig. S1B). Coincidently, we have previously identified the *smu995-998* operon as the site of suppressor mutations that reverted the hypersensitivity of a Δ*dpr* strain to oxidative stresses (20). Notably, Dpr is an iron-sequestering intracellular protein that plays a major role in bacterial oxidative stress tolerance (21). While a mechanistic explanation for the association of the Smu995-998 operon with restoration of arginine tolerance in the Δ*spxA1* is not the focus of this report, we predict that reductions in intracellular iron pools in the suppressor mutants reduced Fenton reactivity at elevated pH, which compensates for the loss of SpxA1-dependent antioxidant activity.

In addition to SNPs within the *smu995-998* operon, all sequenced isolates, including the original arginine-sensitive Δ*spxA1* isolate, bore a single base deletion at position 393 within the coding region of the *smu593* gene (Fig. 1A), which is predicted to code for the peroxide regulator PerR. This single base deletion introduced a frameshift mutation that resulted in a premature truncation of the PerR protein (Fig. 1B). In light of this unexpected finding, we used PCR sequencing to determine if this *perR* mutation was present in our freezer stocks of strain UA159 and Δ*spxA1*. To our surprise, the frameshift mutation discovered in Δ*spxA1* was found in our original UA159 freezer stocks. We contacted six other labs that routinely work with *S. mutans* UA159 and found that UA159 stocks from some of these labs had the same single base deletion at position 393 while one other lab had a PerR C106F codon substitution mutation. We suspect that the *perR* mutation at position 393 occurred early on and was passed among different labs over the years. Nonetheless, the *perR* gene sequence in stocks from two labs was intact as well as in a lyophilized stock that was purchased from ATCC.

**Fig. 1.**
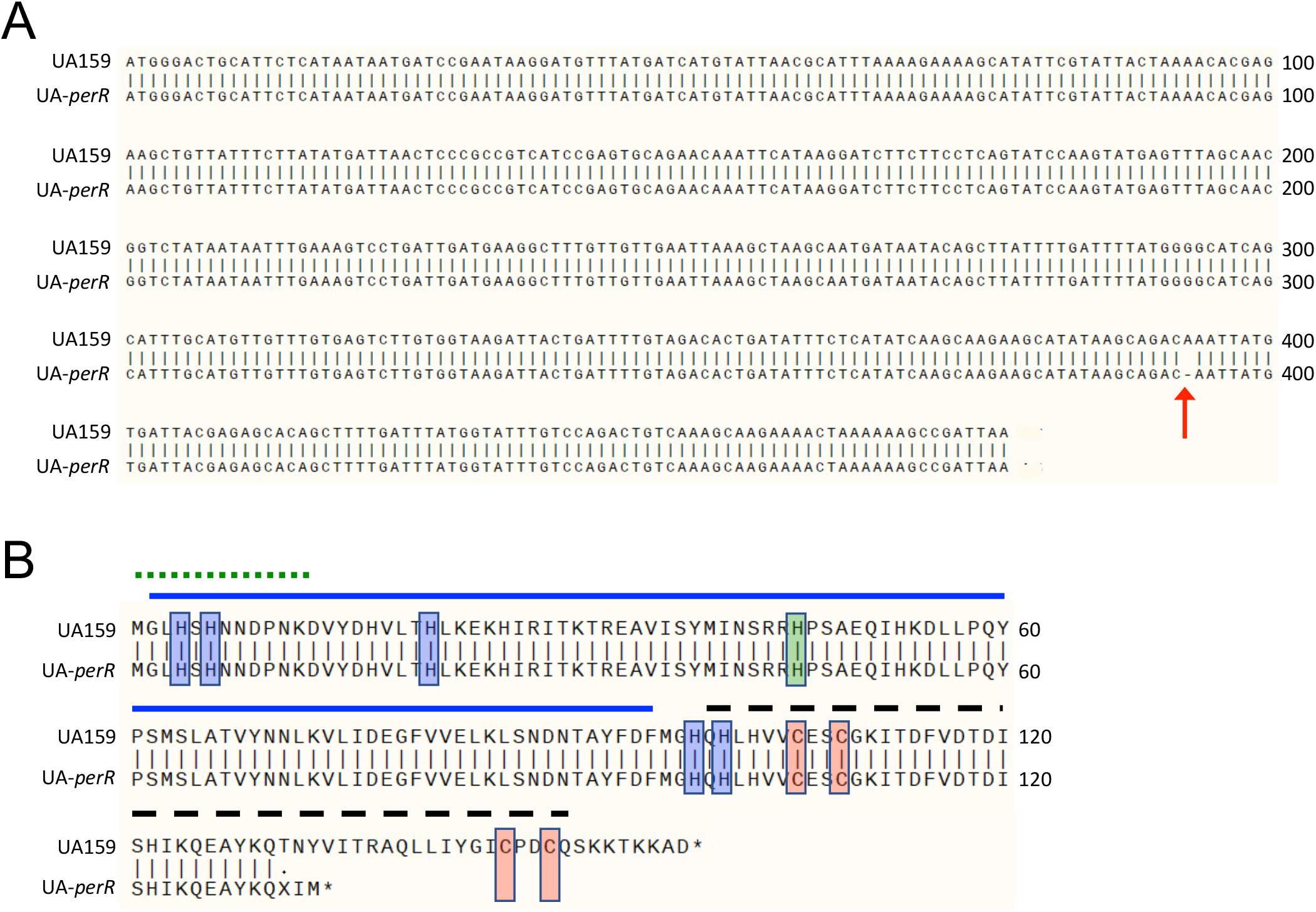
Whole genomic sequencing reveals a *perR* SNP that results in a prematurely truncated protein in *S. mutans* UA159. (A) Alignment of nucleotide sequences of *perR* from UA159 from the NCBI (*perR*+) and UA-*perR* (*perR*−). The red arrow at base 393 indicates a 1-bp deletion in the sequence of UA-*perR*. (B) Alignment of amino acid sequences of PerR from UA159 and UA-*perR*, indicating a premature termination of the latter. Red shading indicates zinc-coordinating cysteine residues critical for PerR function. Blue shading indicates histidine residues involved in binding of a regulatory Fe molecule. Green shading indicates a histidine residue believed to be important in DNA binding. The solid blue line indicates the N-terminal DNA-binding domain. The dashed black line indicates the C-terminal dimerization domain. The dotted green line indicates an N-terminal extension unique to PerR members of the Fur family and conserved among streptococci. Structural domains were based on homologies to those described in PerR of *S. pyogenes*, whose crystal structure has been solved (22).

Similar to most members of the Fur family, the PerR proteins of *B. subtilis* (PerR_BS_) and *S. pyogenes* (PerR_SP_) require a zinc molecule for structural stability (8, 22). This zinc is tightly coordinated by cysteines that compose two highly conserved CXXC redox switch domains found in each member of the Fur family. The zinc molecule that is bound cooperatively by these 4 cysteine residues contributes to the structural stability and to the capacity for dimerization of PerR (23). The premature *perR* stop codon identified in some of the UA159 stocks indicates that the second CXXC motif is missing in the truncated PerR protein while the C106F codon substitution in the UA159 stock from one other lab directly disrupts the first CXXC motif. Thus, both the frameshift mutation at codon 393 and C106F amino acid substitution are predicted to affect the structure of the zinc-binding domain that constitutes part of the dimer interface of PerR. For the remaining parts of this manuscript, we renamed our UA159 isolate containing the truncated PerR as UA-*perR*, maintaining the UA159 designation for the strain harboring the intact *perR* gene.

Next, we used the intact UA159 PerR amino acid sequence to BLAST against sequenced *S. mutans* genomes. Out of the dozens of *S. mutans* PerR sequences currently available at the NCBI, the overwhelming majority were found to have no important changes at the amino acid level as compared to the original UA159 sequence. We also amplified and sequenced the *S. mutans perR* open reading frame from several fresh clinical isolates available in the lab and found that the *perR* gene was intact in all these strains (data not shown). Interestingly, we found that two other “laboratory strains”, NG8 and GS-5, also displayed *perR* SNPs. The GS-5 strain displayed a A36D amino acid substitution while the NG8 strain carried a 7-bp deletion that resulted in a truncation of the PerR protein even more premature than that of our UA159 stock (Fig. S2). Based on this analysis, we conclude that the PerR protein sequence is fairly conserved in *S. mutans*, though possibly a genetic hotspot of spontaneous mutations in laboratory strains.

### Disruption of PerR enhances the tolerance of *S. mutans* to peroxide stress

Next, we sought to determine the impact of the PerR truncation in our UA159 stock (UA-*perR*), and more broadly the importance of PerR to *S. mutans* physiology by replacing the entire *perR* gene in UA159 with an erythromycin resistance cassette to generate the UAΔ*perR* strain. When grown in a 5% CO_2_ atmosphere using BHI, UAΔ*perR* and UA-*perR* displayed a slight growth advantage as compared to UA159 of the complemented UAΔ*perR* (Fig. S3). Next, we grew the strains in BHI at 37°C in an automated growth reader containing an oil overlay on top of each well to create a semi-anaerobic environment. When grown in BHI alone in this environment, UA-*perR* displayed a slight growth advantage compared to UA159 and UAΔ*perR* (Fig. 2A) similar to the differences observed in the 5% CO_2_ atmosphere (Fig. S3). When grown in media containing 0.4 mM H_2_O_2_, the UA159 strain displayed an extended lag (~ 10 hours) while UA-*perR* and UAΔ*perR* showed a much shorter adaptation phase (< 4 hours) (Fig. 2B). In addition, UA159 was completely unable to grow in BHI containing 0.5 mM H_2_O_2_ whereas both UA-*perR* and UAΔ*perR* grew after an extended lag phase (> 8 hours, data not shown). Because peroxide stress and iron homeostasis are intertwined due to their role in hydroxyl radical generation via Fenton chemistry, we also tested the sensitivity of the different strains to streptonigrin, an iron-dependent antibiotic (24) used as a proxy to determine intracellular iron availability. Here, both UA-*perR* and UAΔ*perR* grew significantly better than UA159 reaching a significantly higher final growth yield, indicating that PerR also controls intracellular iron homeostasis (Fig. 2C). This possibility can be further supported by evidence that PerR controls expression of iron and manganese transporters of other streptococcal species (7, 25, 26). To further investigate this connection, we compared intracellular iron and manganese content of BHI-grown UA159, UAΔ*perR* and UA-*perR* strains. However, total intracellular metal quantifications revealed similar profiles for intracellular iron and manganese among the different strains (Fig. S4A), suggesting that the streptonigrin sensitivity of PerR-negative strains might be due to differences in freely-available iron. To investigate this possibility, we took advantage of the availability of an anti-Dpr antibody (20) to determine whether the Dpr protein was more abundant in strains lacking a functional PerR. Indeed, we observed greater quantities of Dpr protein in UA-*perR* and in UAΔ*perR* as compared to the UA159 and UAΔ*perR*^Comp^ strains (Fig. S4B). In streptococci, Dpr is the primary iron-binding protein within the cell (21), and its ability to sequester iron molecules minimizes the generation of ROS through Fenton chemistry. Thus, the increased abundance of Dpr in *perR*-defective strains may help explain, at least in part, why the UA-*perR* and UAΔ*perR* strains were more tolerant to streptonigrin despite showing similar intracellular iron concentrations when compared to PerR^+^ strains.

**Fig. 2.**
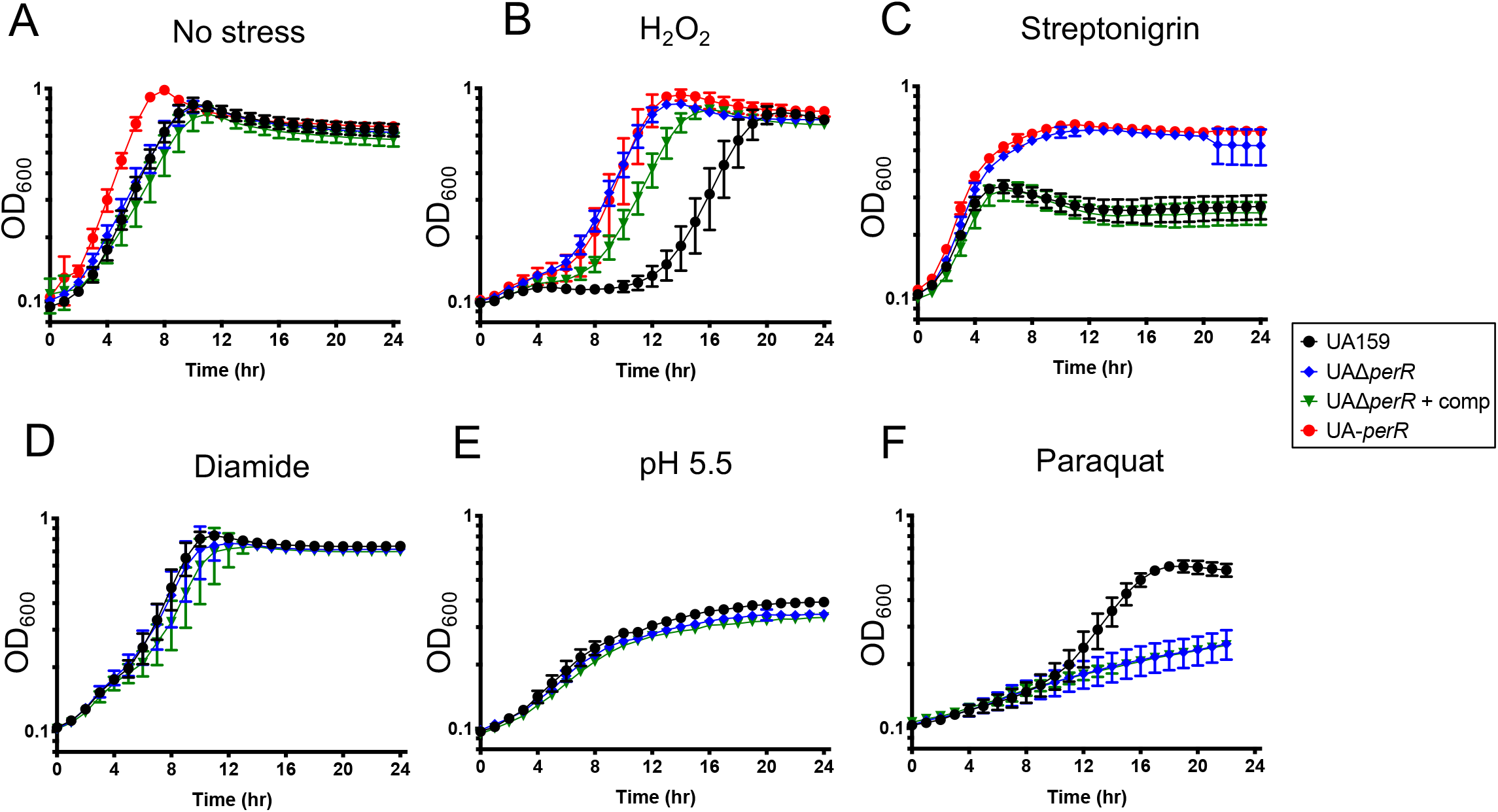
Disruption of *perR* enhances growth of *S. mutans* in the presence of stresses. Growth of UA159 (*perR*+), UA-*perR* (*perR*−), the *perR* deletion mutant strain (UAΔ*perR*) and complemented strain (UAΔ*perR*^Comp^). Strains were grown in (A) BHI, (B) BHI containing 0.4 mM H_2_O_2_, (C) BHI containing 0.2 μg ml^-1^ streptonigrin, (D) BHI containing 0.8 mM diamide, (E) BHI adjusted to pH 5.5, or (F) BHI containing 10 mM paraquat. Data represent averages and standard deviations of results from four independent experiments.

Given the phenotypic similarities of UA-*perR* and UAΔ*perR*, we opted to continue our investigations only with UAΔ*perR* and its genetic complemented version. Different than H_2_O_2_ or streptonigrin, a growth advantage for UAΔ*perR* was not observed in the presence of the thiol-stressor diamide (Fig. 2D), or at low pH conditions (Fig. 2E). Moreover, Δ*perR* was more sensitive to growth in the presence of paraquat, a superoxide generating compound, when compared to the parent UA159 strain (Fig. 2F). These observations are in line with a previous report showing that *perR* inactivation enhanced peroxide tolerance of *S. mutans* (27) and of closely-related species such as *Streptococcus pyogenes* and *Enterococcus faecalis* (28–30) while increasing the superoxide stress sensitivity of *S. pyogenes* (10, 22, 28, 29, 31). Thus, the relationship of PerR regulation, oxidative stress and metal homeostasis appears to be rather complex with our results reinforcing that PerR has a primary role in (if not restricted to) the peroxide stress response. The phenotypes of the Δ*perR* strain were fully (Fig. 2A and C) or partially (Fig. 2B) restored in the genetically-complemented UAΔ*perR*^Comp^ strain. Given that the genetic complementation was obtained by integrating a single copy of *perR* elsewhere in the chromosome, the reason for the partial complementation in the presence of H_2_O_2_ is unknown.

While the premature protein truncation in *S. mutans* NG8 almost certainly rendered PerR inactive, the consequence of the PerR A36D amino acid substitution in the GS-5 strain is unclear (Fig. S2). To begin to understand the consequence of these mutations to each respective strain, we transformed NG8 and GS5 with the pMC340B-*perR* integration plasmid that was used to complement UAΔ*perR*. Genetic complementation of *perR* in NG8 and GS-5 did not affect growth kinetics in BHI medium; however, both complemented strains became more sensitive to peroxide stress showing an extended lag phase when grown in media containing 0.3 mM H_2_O_2_ (Fig. 3). Thus, it appears that the emergence of *perR* loss-of-function mutations may not be an uncommon phenomenon in laboratory strains of *S. mutans*.

**Fig. 3.**
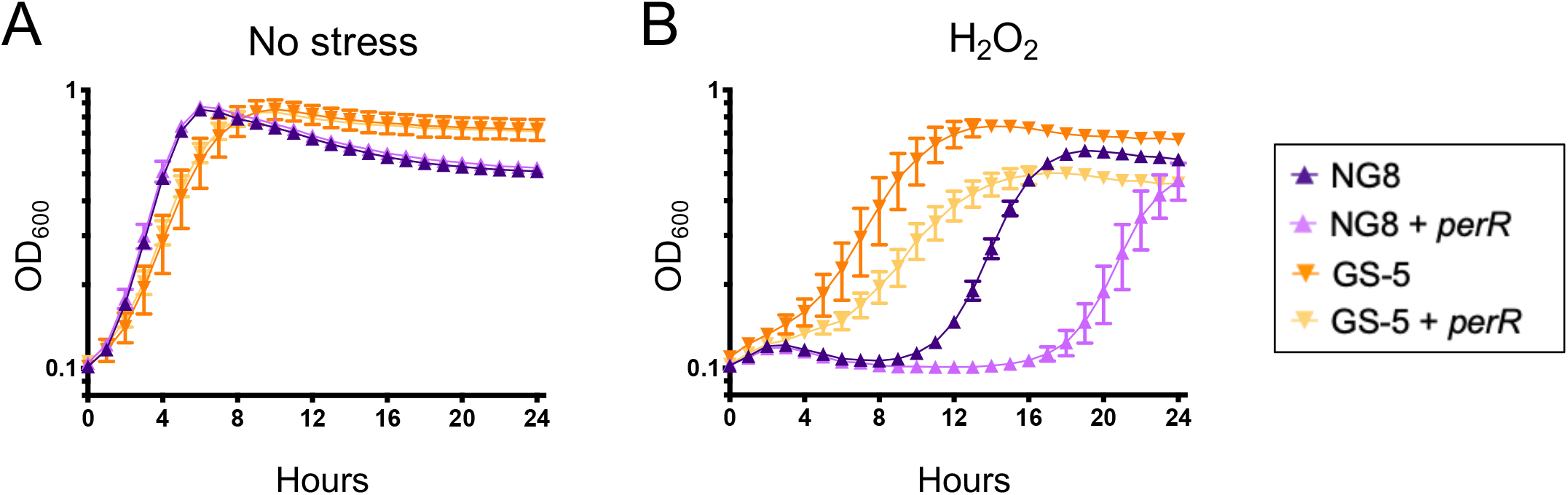
Chromosomal integration of an intact copy of the *perR* gene diminishes H_2_O_2_ tolerance in *S. mutans* NG8 or GS-5. Growth curves in (A) BHI or (B) BHI containing 0.3 mM H_2_O_2_. Data represent averages and standard deviations of results from four independent experiments.

### Exploration of the regulatory reach of PerR

Based on extensive evidence that PerR functions as a peroxide stress regulator in gram-positive bacteria (4, 28–30), including *S. mutans* (27), we next used RNAseq to determine the scope of the PerR regulon. As UA-*perR* and UA159 Δ*perR* showed similar phenotypes, we chose to compare the transcriptome of UAΔ*perR* to that of UA159. Briefly, UA159 and UAΔ*perR* were grown to mid-exponential phase and exposed to 0.5 mM H_2_O_2_ for 5 minutes, a condition that we had previously demonstrated as optimal for activation of the peroxide stress response (19). As we have shown before (19), peroxide stress resulted in a strong and rapid induction of SpxA1-regulated genes including archetypal oxidative stress genes such as *ahpCF, dpr, tpx* and *sodA* (Table S1). Surprisingly, the peroxide stress transcriptome of UAΔ*perR* (data not shown) was nearly identical to the transcriptional profile of UA159 subjected to H_2_O_2_ stress. Yet, when directly compared to UA159, 40 genes (4 upregulated, 36 downregulated) in the unstressed condition and 60 genes (19 upregulated, 41 downregulated) following exposure to H_2_O_2_ were differentially expressed in the UAΔ*perR* strain (False Discovery Rate 0.05, 2-fold cutoff, Table S2). Notably, all genes differentially expressed by UAΔ*perR* in the unstressed condition showed the same expression trends following H_2_O_2_ exposure. Unexpectedly, none of the typical PerR-regulated genes (e.g. *dpr, tpx* and *sodA*) were differentially expressed in the UAΔ*perR* strain when compared to UA159 under any given growth condition. A number of the differentially expressed genes belong to two large polycistronic transcriptional units (*smu193c-smu217c*) and (*smu1750c-1764c*) coding, respectively, for hypothetical proteins of unknown function and the CRISPR2-*cas* system, a defense mechanism believed to provide immunity against phage invasion (32). In *S. mutans* UA159, the two CRISPR systems are also linked to stress responses, though the reason for this association remains to be determined (33). For visualization purposes, a dot plot graph showing all genes differently expressed in UAΔ*perR* is provided (Fig. 4). It is worth noting that the changes in gene expression in most cases were rather modest as can be visualized by the large number of genes clustered closely below or above the x-axis. While we expected that at least a subset of oxidative stress genes would be differently expressed in the Δ*perR* strain, a previous transcriptome analysis of an *S. pyogenes* Δ*perR* strain revealed that only 6 genes were strongly regulated by PerR, with only the iron efflux *pmtA* gene later confirmed to be under direct PerR control (7). Also in *S. pyogenes*, PerR was found to bind specifically to the *ahpC* promoter region but transcriptional analysis failed to confirm a role for PerR in *ahpC* transcriptional expression (31). Finally, PerR was also shown to exert fairly modest transcriptional control of a small number of genes in *E. faecalis* (30). Collectively, these observations indicate that PerR does not exert major transcriptional control over the oxidative stress genes of these bacteria supporting previous evidence that activation of the peroxide stress response of *S. mutans* is primarily controlled by SpxA1 (19).

Because global transcriptional approaches are not as sensitive as targeted approaches and may overlook small, yet meaningful, changes in transcription, we also used real time quantitative PCR (RT-qPCR) to compare the expression of a subset of oxidative stress genes, with or without putative PerR-binding DNA motifs, in the UA159, UA-*perR*, UAΔ*perR* and UAΔ*perR*^Comp^ strains. Starting with the premise that SpxA1-mediated activation during peroxide stress overrules PerR-mediated effects on genes controlled by both regulators, this analysis was focused on cells grown in the absence of stress. PerR is reported to bind to a 15-bp palindrome motif (TTANAATNATTNTAA) within the promoter region of its targeted genes (31). This motif is similar to that recognized by Fur, maintaining the 7-1-7 arrangement though it has been noted that a number of PerR-regulated genes do not have perfect repeats (6). Despite the similarity of PerR and Fur operators, there is little overlap of these regulators and their targets (12). For example, Fur binding sites can be distinguished from PerR boxes by guanine and cytosine residues that are recognized by a conserved arginine residue present in the Fur protein but not in PerR (34). To identify putative PerR-regulated genes in the *S. mutans* genome, we accessed the RegPrecise database (35) and manually scanned the promoter regions of selected oxidative stress and metal transport genes. RegPrecise listed five putative PerR-regulated genes (*feoA*, *dpr, sloA, smu635*, and *perR* itself) that contained one or two mismatches when compared to the consensus sequence (Table 1). Our manual searches identified four additional genes (*mntH*, *spxA1*, *sodA* and *smu995*) with putative PerR-binding boxes albeit, these additional motifs contained four mismatches from the consensus, with the exception of *sodA* that contained only one mismatch (Table 1). Of note, we recently showed that *spxA1* is co-repressed by PerR and SloR in a redundant manner as only the simultaneous inactivation of *perR* and *sloR* increased *spxA1* transcription (36). Overall, a trend was observed showing that expression levels of most of the genes tested were elevated in UA-*perR* and UAΔ*perR* as compared to UA159, the iron transport systems *feoAB* and *smu995-998* were the notable exceptions (Fig. 5). Interestingly, *smu635* coding for a hypothetical protein of unknown function, seemingly unique to the streptococci, was the most upregulated gene in the absence of PerR. Work is currently underway to determine the function of Smu635.

**Fig. 4.**
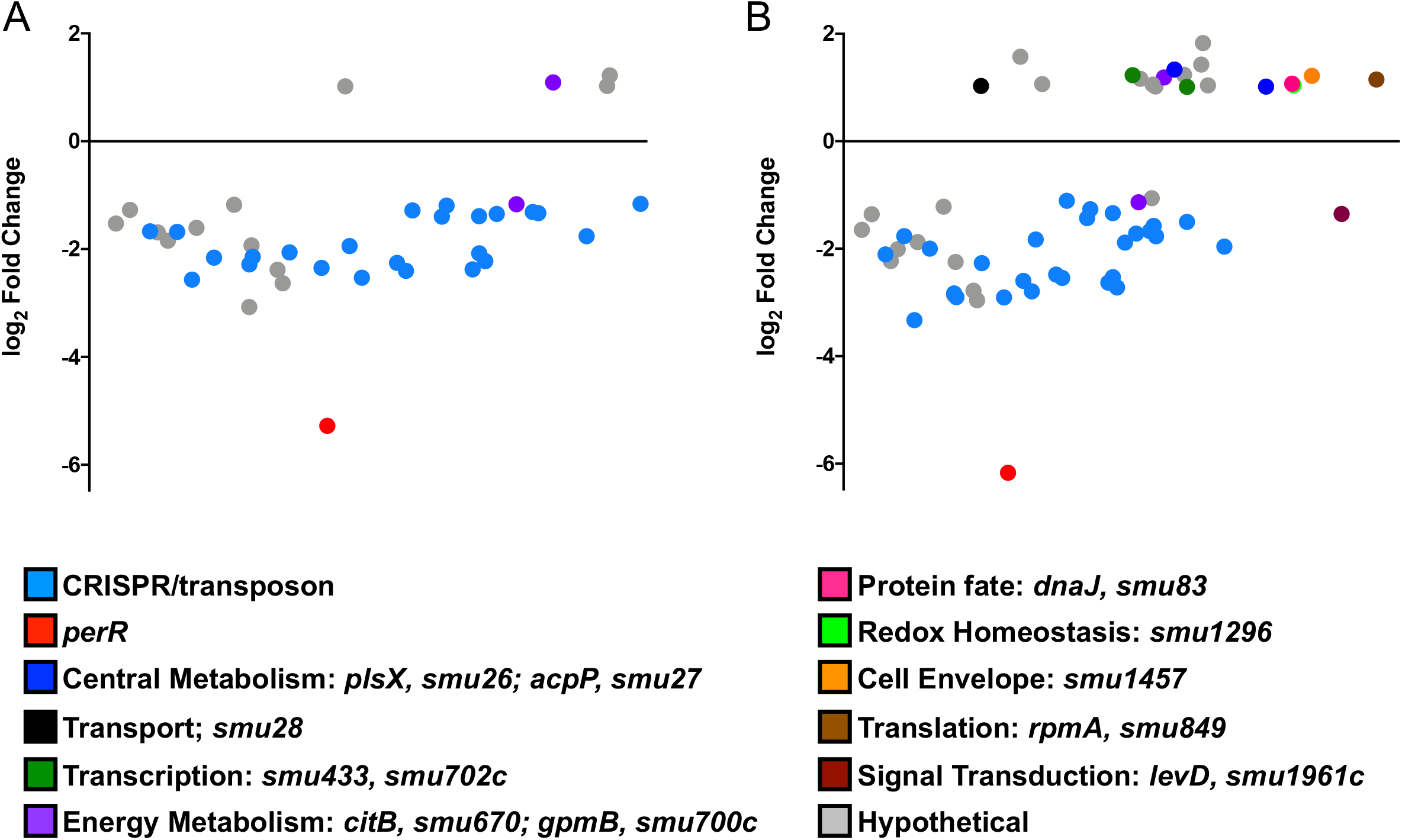
Dot plot of genes differentially expressed in UAΔ *perR* as compared to UA159 as determined by Degust (degust.erc.monash.edu) under (A) control or (B) H_2_O_2_ stress conditions. The y-axis indicates the log_2_ fold change in expression while the x-axis indicates the average expression level of each gene compared to all other genes. The details of the expression trends are shown in Table S2.

**Fig. 5.**
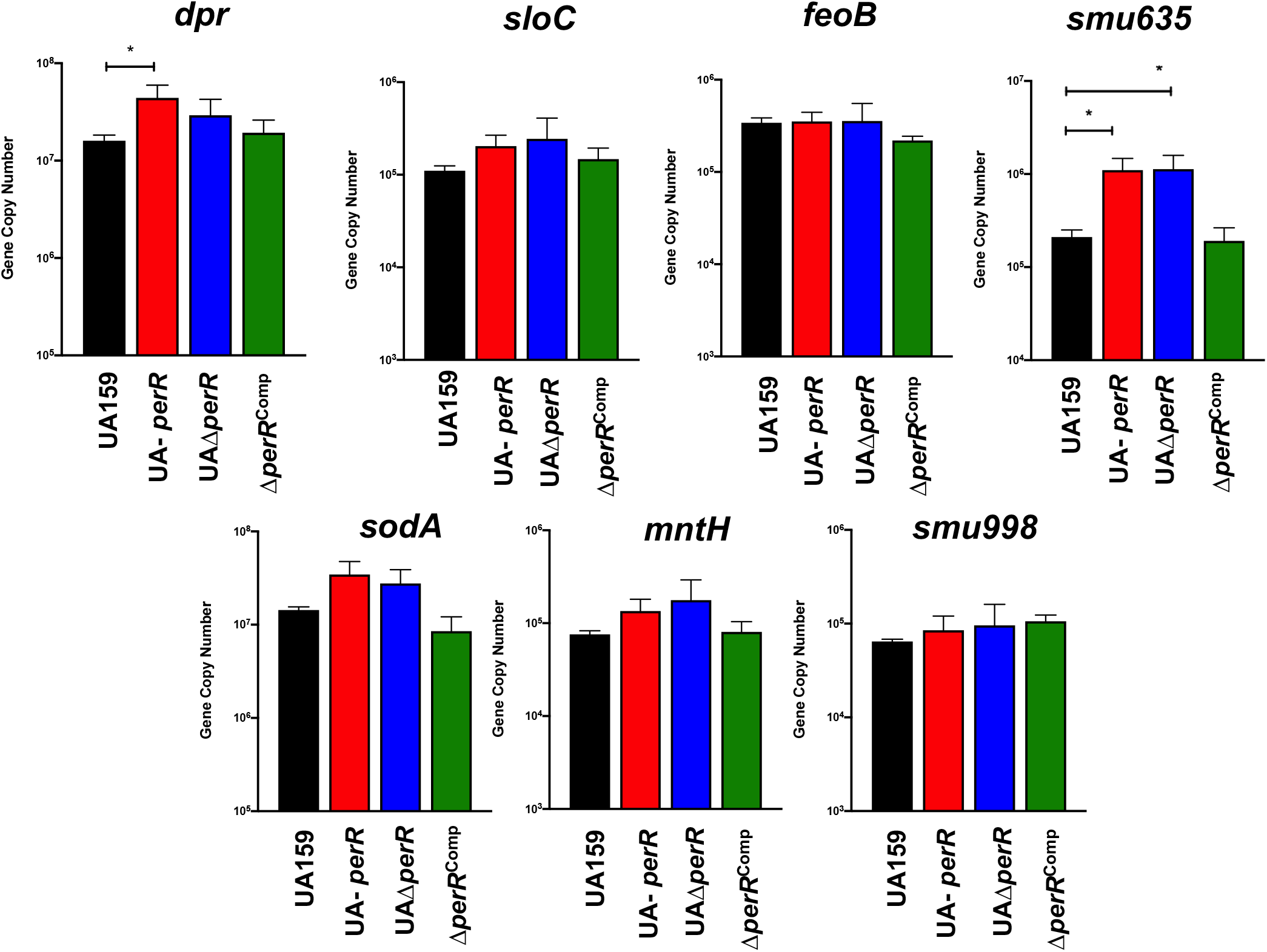
RT-qPCR analysis of genes with putative PerR-binding motifs reveals a trend of PerR transcriptional repression. Bars represent the gene copy number and represent averages and SDs from three replicate samples. ANOVA was performed to verify significance of the data; expression levels of each gene was compared to that of UA159. (*, *p* ≤ 0.03)

**Table 1.**
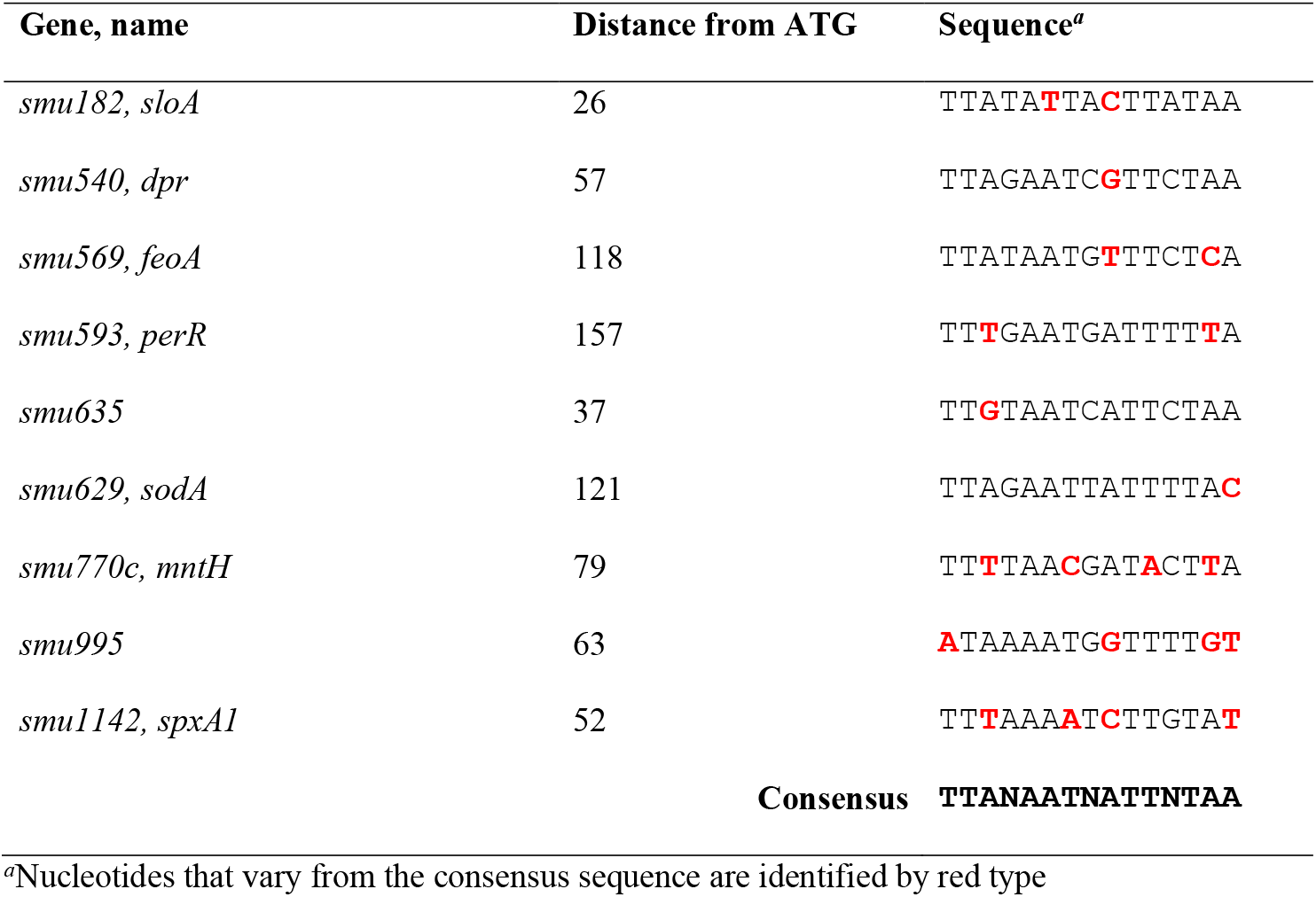
Putative PerR-binding motifs in the *S. mutans* UA159 genome.

### Concluding remarks

The accidental discovery of spontaneously-occurring *perR* gene mutations in laboratory stocks of UA159, the first fully sequenced and (by far) the most thoroughly characterized lab strain of *S. mutans*, supports anecdotal observations of bacterial adaptation to *in vitro* growth conditions. As shown in this report, loss of PerR regulation conferred a growth advantage to *S. mutans* under conditions (i.e., in the presence of air) that are routinely encountered in the laboratory setting. Interestingly, loss-of-function *perR* mutations were also identified in strains GS-5 and NG8, two of the most commonly used laboratory strains after the UA159 strain, suggesting that routine conditions used to cultivate *S. mutans* in the laboratory may facilitate the emergence of *perR* inactive strains. However, how the loss of PerR regulation increased peroxide tolerance without making a significant impact in oxidative stress gene expression remains to be understood.

It has been proposed that tight regulation of stress genes is important because expression of these genes in the absence of stress may decrease overall fitness (37, 38). In both *S. pyogenes* and *E. faecalis*, the ability of *perR* mutant strains to tolerate peroxide stress *in vitro* did not correlate with the ability of those strains to cause disease (10, 30, 31), indicating that PerR is required to fine-tune gene expression during infection. Thus, the growth benefit acquired by *S. mutans perR* mutants under *in vitro* conditions is unlikely to arise during host colonization providing a possible explanation as to why *S. mutans* have retained a functional *perR*. Studies to determine the significance of *S. mutans perR* during host colonization and disease progression are still warranted.

This report must also serve as a cautionary tale for conclusions broadly based on a single bacterial strain and the importance of keeping culture passages to a bare minimum, and of routine (re)assessment of strains’ genetic and phenotypic traits. While these precautions may not completely eliminate the emergence of spontaneous mutants, especially those that confer a growth advantage, the accessibility and cost reductions of WGS technologies have made identification of these occurrences a manageable undertaking.

## MATERIALS AND METHODS

### Bacterial strains and growth conditions

The bacterial strains used in this study are listed in Table 2. Strains of *S. mutans* were routinely grown in brain heart infusion (BHI) at 37°C under anaerobic conditions. For physiologic analysis, overnight cultures were sub-cultured 1:20 in BHI medium in a 5% CO_2_ atmosphere and growth was monitored over time. To evaluate the ability of the different strains to grow in the presence of stress conditions, cultures were grown to an optical density at 600nm of 0.25, and then diluted 1:50 into the appropriate medium. Growth was then monitored in a microtiter plate with an overlay of sterile mineral oil to minimize oxygen exposure using an automated growth reader (Bioscreen C) at 37°C. For growth in the presence of arginine, overnight cultures were serially diluted and 10 μL spotted onto TYG agar with or without arginine supplementation. For gene or protein expression analysis, overnight cultures were subcultured 1:20 in BHI as described above and grown to an optical density at 600nm of 0.4, at which point control samples were harvested by centrifugation. For RNA isolation, harvested pellets were resuspended in 1 ml RNA Protect Bacteria Reagent (Qiagen), incubated for 5 min at ambient temperature, harvested by centrifugation and the pellets stored at −80°C until use.

**Table 2.** Bacterial strains used in this study.

| Strain | Relevant characteristics | Source |
| --- | --- | --- |
| *S. mutans* UA159 | Wild-type | ATCC |
| *S. mutans* UA-*perR* | UA159 variant with *perR* mutation | Spontaneously occurring/<br>Laboratory stock |
| *S. mutans* Δ*spxA1* | *spxA1*::Spec | (42) |
| *S. mutans* UAΔ*perR* | *perR*::Erm | This study |
| *S. mutans* UAΔ*perR* + *perR* | *perR*::Erm, *mtl*-, *perR*+ | This study |
| *S. mutans* NG8 | Wild-type, truncated PerR | Laboratory stock |
| *S. mutans* NG8 + *perR* | *mtl*-, *perR*+ | This study |
| *S. mutans* GS-5 | Wild-type, PerR<sup>A36D</sup> | Laboratory stock |
| *S. mutans* GS-5+ *perR* | *mtl*-, *perR*+ | This study |

### DNA purification for whole genomic sequencing

For DNA isolation, cells from 25 ml stationary phase cultures of *S. mutans* were harvested by centrifugation (3000 x g, 15 min), suspended in a mixture of 50 mM NaCl, 25 mM EDTA and lysozyme (1 mg ml^-1^) and incubated for 3 hours at 37°C. Then, 0.1 mm glass beads and 1% sodium lauryl sulfate (SDS) were added to the suspension, which was subjected to high speed vortexing for 5 min, followed by an additional 30 min incubation at 37°C. Nucleic acid was extracted with 1 volume of buffer-saturated phenol. The aqueous phase was collected and extracted with phenol:chloroform:isoamyl alcohol followed by another centrifugation step. The aqueous phase was precipitated with 0.3 M sodium acetate pH 5.5 and an equal volume of cold absolute ethanol. The precipitated DNA was washed with 70% ethanol, and then resuspended in sterile deionized H2O containing RNAse. After 30 min incubation at room temperature, the DNA solution was again extracted with phenol:chloroform:isoamyl alcohol and ethanol precipitated as described above. The DNA was examined by agarose gel electrophoresis to ensure that the preparation contained high molecular weight DNA free of RNA. The DNA was used for WGS with SNPs and indels detected using CLC Genomics Workbench at Tufts University Core Facility (Boston, MA) using *S. mutans* UA159 (NC_004350) as the reference sequence.

### Construction of mutant and complemented strains

A UA159 (*perR*^+^) strain lacking the *perR* gene was constructed using a PCR ligation mutagenesis approach (39). Briefly, PCR fragments flanking the *perR* gene were ligated to an erythromycin resistance cassette after digestion with the same restriction enzymes. This ligation mixture was used to transform *S. mutans* UA159; the recombinant strain was isolated on BHI agar supplemented with 10 μg ml^-1^ erythromycin. The gene deletion was confirmed by sequencing the amplicon containing the antibiotic cassette insertion site and flanking region. The mutant strain was complemented by cloning the full-length *perR* gene (and native promoter) into the *S. mutans* integration vector pMC340B (40) to yield plasmid pMC340B-*perR*. The plasmid was used to transform the *S. mutans ΔperR* strain for integration at the *mtl* locus. This same strategy was also used to transform *S. mutans* GS-5 and NG8 strains. All primers used in this study are listed in Table 3.

**Table 3.**
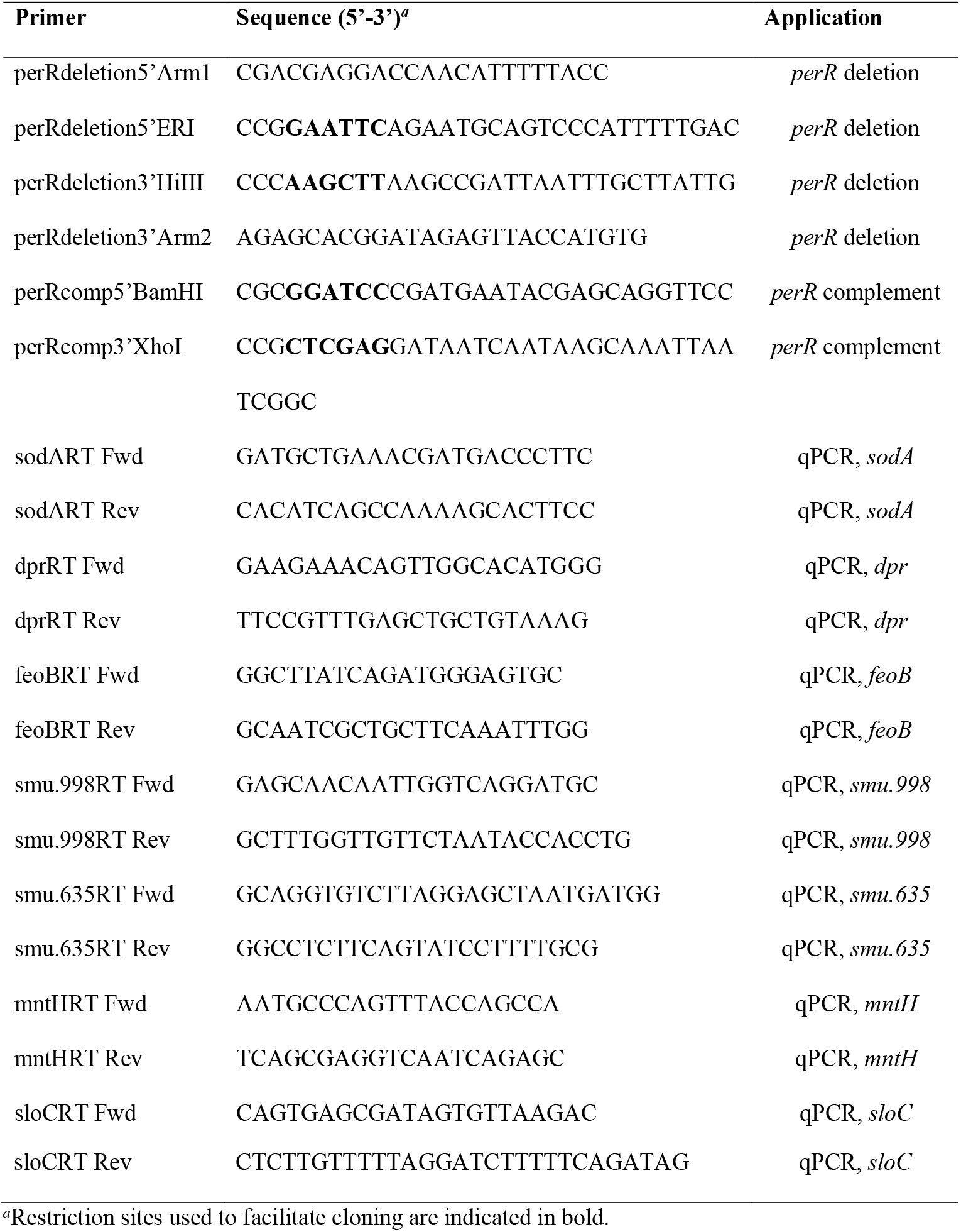
Primers used in this study.

### ICP-MS analysis

The total metal content within bacterial cells was determined using Inductively Coupled Plasma Mass Spectrometry (ICP-MS) performed at the University of Florida Analytical Toxicology Core Laboratory. Briefly, cultures were grown to mid-exponential phase (OD_600_ = 0.4) in BHI, harvested by centrifugation at 4°C for 15 min at 4,000 rpm, and washed first in phosphate-buffered saline (PBS) supplemented with 0.1 mM EDTA to chelate extracellular divalent cations followed by a wash in PBS alone. Cell pellets were resuspended in HNO3 and metal composition was quantified using a 7900 ICP mass spectrometer (Agilent). Metal concentrations were then normalized to total protein content determined by the bicinchoninic acid (BCA) assay (Pierce).

### Western blot analysis

Whole-cell protein lysates of mid-log grown cultures of *S. mutans* UA159, UA-*perR*, UAΔ*perR*, or UAΔ*perR*^Comp^ strains were obtained by homogenization in the presence of 0.1 mm glass beads using a bead beater. The protein concentration was determined using the BCA assay. Equal amounts of the protein lysates were separated by 13.5% SDS-PAGE and transferred to a polyvinylidene fluoride (PVDF) membrane (Prometheus). Dpr detection was performed using rabbit anti-rDpr polyclonal antibody (20) diluted 1:1000 in phosphate-buffered saline (PBS) containing 0.05% Tween 20 (PBS-T) followed by anti-rabbit horseradish peroxidase (HRP)-coupled antibody (Sigma-Aldrich) diluted 1:1000 in PBS-T. Immune reactivity was visualized by incubation of the blot with 3,3-diaminobenzidine (Sigma-Aldrich) substrate solution for horseradish peroxidase conjugates. As a loading control, duplicate gels were stained with SYPRO Ruby protein gel stain (ThermoFisher Scientific) and visualized with UV light.

### RNA analysis

Total RNA was isolated by acid-phenol chloroform extraction method from homogenized cell lysates that was modified from a previously described protocol (41). Briefly, harvested cells were resuspended in Tris-EDTA (50 mM:10 mM) in the presence of acid phenol:chloroform (5:1) and SDS detergent, mixed with 0.1 mm glass beads and subjected to homogenization in a Bead beater (BioSpec Products) for three cycles of 30 sec, with 3 min on ice between cycles. The cell lysates were subjected to centrifugation (14000 x g, 10 min) and the upper aqueous phase containing nucleic acid transferred to a new tube. An equal volume of acid phenol:chloroform was vigorously mixed with the aqueous phase by vortexing, followed by incubation for 5 min at ambient temperature, and the upper aqueous phase collected after another centrifugation cycle. This nucleic acid-containing aqueous phase was digested with Ambion DNase I (ThermoFisher) according to the manufacturer’s instructions. Then, RNA was purified using an RNeasy kit (Qiagen) including an on-column DNase digestion according to the manufacturer’s instructions. The final RNA concentration was determined by measuring on a NanoDrop One Spectrophotometer (ThermoFisher). Quantifications of mRNA were obtained by reverse transcriptase quantitative real-time PCR (RT-qPCR) according to protocols described elsewhere (41) using gene-specific primers listed in Table 3. ANOVA was performed to verify significance of the RT-qPCR results using Dunnett’s test to compare expression to that of *S. mutans* UA159.

For RNA-Seq analysis, samples were prepared as previously described (33). Sample quality and quantity were assessed on an Agilent 2100 Bioanalyzer at the University of Florida Interdisciplinary Center for Biotechnology research (UF-ICBR). RNA (5 μg per sample) was subjected to two rounds of mRNA enrichment using a MICROBExpress bacterial mRNA purification kit (Thermo Fisher). cDNA libraries with unique barcodes were generated from 100 ng enriched mRNA using NEB Next UltraII Directional RNA Library Prep kit for Illumina (New England Biolabs), then assessed for quality and quantity by Qubit. Pooled cDNA libraries were subjected to RNA deep sequencing (RNA-Seq) using the Illumina NextSeq 500 platform. Read mapping was performed on a Galaxy server hosted by the UF Research Computer using Map with Bowtie for Illumina and the *S. mutans* UA159 genome (GenBank accession no. NC_004350.2) as a reference. The reads per open reading frame were tabulated with htseq-count. Final comparisons were performed with Degust (http://degust.erc.monash.edu/), with a false-discovery rate (FDR) of 0.05 and a 1.5-fold change cutoff.

### Data availability

Gene expression data have been deposited in the NCBI Gene Expression Omnibus(GEO) database (https://www.ncbi.nlm.nih.gov/geo) under GEO Series accession number GSE158080.

## Acknowledgements

This study was supported by NIH-NIDCR award DE019783. We thank our colleagues from the “*Strep. mutans* community” for sharing strains and helpful discussions.

